# Association of Arg389Gly β1-adrenergic receptor polymorphism with effective dose of β blocker in congestive heart failure among Chinese Han population

**DOI:** 10.1101/272013

**Authors:** Rong Gu, Yu Shen, Jianhua Liu, Shuaihua Qiao, Lian Wang, Biao Xu

**Author notes:** **Corresponding authors at** Department of Cardiology, Affiliated Drum Tower Hospital, Nanjing University Medical School, Zhongshan Road, Nanjing 210008, China.

## Abstract

**Background:** To investigate the relationship between β1-adrenergic receptor (ADRB1) gene polymorphisms and the response to medication in patients with congestive heart failure (CHF).

**Methods and Results:** Two hundred and sixty patients with CHF were enrolled. Ser49Gly and Arg389Gly polymorphisms were identified. During one-year follow-up, differences of echocardiographic parameters and major adverse cardiac events (MACE) were analyzed. For Ser49Gly polymorphisms, there were no differences between AA genotype and AG/GG genotype of baseline clinical features and echocardiographic parameters as well as one-year incidence of MACE. For Arg389Gly polymorphisms, there were no significant differences in baseline clinical characteristics, LVDd and LVEF among the three genotypes. However, the increase amplitude of LVEF after one year among patients carrying GG genotype was significantly higher than those carrying CC genotype (11.7% vs 1.3%, *P*<0.05). The incidence of MACE among different genotypes of CC, CG and GG were 22.2%, 10.0% and 8.3%,with statistical difference (*P*=0.021).

**Conclusions:** The study suggested there was no relationship between Ser49Gly polymorphisms of the ADRB1 gene and the therapeutic effect and prognosis in CHF patients under the same dosage of drugs. However, the improvement of cardiac function and prognosis in patients carrying the Gly389 allele were significantly better than those of Arg389Arg homozygous.

## 1. Introduction

Congestive heart failure (CHF) is a complex clinical syndrome recognized as the end-stage of various cardiovascular diseases. Intensive researches have stated the excessive long-term hyperactivation of the adrenergic nervous system in CHF contributes to disease progression (1). Blockade of β1-adrenergic receptor (ADRB1), the principal subtype of adrenergic receptor on cardiomyocytes, can improve CHF patients’ prognosis.

However, pharmaceutical effects of ADRB1 blockers vary in CHF patients, which might be explained, at least in part, by genetic differences. The coding block of intron-less ADRB1 gene has two common nonsynonymous single nucleotide polymorphisms (SNPs) at nucleotides 145 and 1165(2). Gene polymorphism(rs1801252) locus at nucleic acid 145(A→G) resulted in substitution of Ser for Gly at position 49, and subsequently lead to a lower baseline activity of adenylate cyclase and activation by isoprenaline(3). Arg389Gly gene polymorphism (rs1801253) locus at position 1165, base G is replaced by C. Arg389 variant predisposes to heart failure by instigating hyperactive signaling programs leading to depressed receptor coupling and ventricular dysfunction(4).

However, so far, the results of different researches in this field were inconsistent, even paradoxical, so there is no definite conclusion to guide clinical applications(5,6). Also, few researches were performed among Chinese people.

Our study aimed to explore the relationship of ADRB1 gene polymorphism of Ser49Gly and Arg389Gly and their corresponding therapeutic response.

## 2. Materials and Methods

### 2.1. Study population

We examined 260 adult patients with CHF, who were referred to Cardiology Department, Nanjing Drum Tower Hospital between October 2013 and April 2015. Inclusion criteria: age between 18 and 80 years; etiology of CHF is either idiopathic dilated cardiomyopathy (IDCM) or ischemic heart disease (IHD) which was diagnosed by coronary angiography; left ventricular ejection fraction (LVEF) ≤40% in echocardiography; New York Heart Association (NYHA) Classification Cardiac Function II-III under proper treatment. Exclusion criteria: acute myocardial infarction; secondor third degree atrioventricular block, sick sinus syndrome, cardiac arrest, allergy to β receptor blocker, asthma, severe kidney or liver dysfunction, malignant tumor, life expectancy <1year and cardiac resynchronization therapy with pacemaker function (CRT-P) or cardiac resynchronization therapy with defibrillator function(CRT-D) implantation within 1year.

### 2.2. Automated DNA sequence analysis

Genomic DNA was extracted from peripheral blood utilizing DP348 kit (Tiangen, China). Polymerase chain reaction (PCR) was performed to amplify 2 fragments encompassing the entire ADRB1 gene. The PCR fragments were then purified and subjected to cycle sequencing by overlapping primers and the ABI PRISM BidDye Terminator Ready Reaction Mix (Applied Biosystems, USA). Geno-typing for Codon 49 and Codon 389 of ADRB1were performed with achieved DNA by TaqMan Assay (Thermo Fisher, USA). For Codon 49 polymorphism of ADRB1, the corresponding base of Ser49Ser was AA, Ser49Gly was AG and Gly49Gly was GG; and the forward primer used was 5’-CCGGGCTTCTGGGGTGTTCC-3’ and the reverse primer was 5’-GGCGAGGTGATGGCGAGGTAGC-3’. For the polymorphism of Codon 389, the corresponding base of Arg389Arg was CC, Arg389Gly was CG and Gly389Gly was GG; and the forward primer was 5’-CGCTCTGCTGGCTGCCCTTCTTCC-3’ and the reverse primer was 5’-TGGGCTTCGAGTTCACCTGCTATC-3’.

### 2.3. Clinical data collection

Patients continued pharmacotherapy in outpatient clinic, and the average follow-up period was 1year. All β-receptor blocker used in this study was metoprolol tablets or metoprolol sustained-release tablets. Aldosterone antagonist used was spirolactone. The use of ACEI or ARB was decided individually by physicians according to patients’ condition. Patients’ baseline data, including age, gender, CHF etiology, hypertension, diabetes, smoking habit, creatinine level, BNP, heart rate, and NYHA Classification were all obtained through electronic medical record. Major adverse cardiac events (MACE) were defined as cardiac death, heart transplantation, malignant arrhythmia, rehospitalization due to CHF. 3-month, 6-month and 1-year follow-up were completed through outpatient clinic, returning visit or telephone follow-up.

### 2.4. Statistical analysis

All statistical analyses were performed with SPSS statistical software (version 20.0, Chicago, USA). Quantitative data were expressed as mean ± SD. One-way ANOVA was performed to compare among three groups; independent sample *t*-test was performed to compare between two groups; rank-sum test was performed to compare NYHA Classification. Enumeration data was compared with chi-squared test.

## 3. Results

### 3.1. Baseline Clinical Characteristics

We recruited 260 CHF patients, among which 163 patients (including 121 male and 42 female) have completed full medical records and 1-year follow-up. Mean age was 62.2 ± 15.4 years; baseline heart rate was 78.5± 14.0 bpm. Among these patients, 125 patients was diagnosed IHD; 79 had a history of hypertension; 42 had diabetes; 63 were smokers. There were no significant differences in gender, CHF etiology, and medical history across different genotypes.

### 3.2. Interaction between Codon 49 genotype and treatment

The analysis results of Codon 49 genotype was shown in figure 1A. As shown in Table 1 and 2, there was no significant difference in baseline characteristics, echocardiographic data and pharmacotherapy during 1-year follow-up period (*P*>0.05). Compared with the G-allele carriers, a homozygote tended to obtain better systolic function recovery and more left ventricular size reduction according to echocardiography within 12-months follow-up; however, they were not statistically different (Figure 2 and Table 3 and 4). 1-year MACE incidence among patients with AA genotype was13.7%; whereas, it was 21.4% among G-allele carriers (*P* = 0.208).

**Figure 1.**
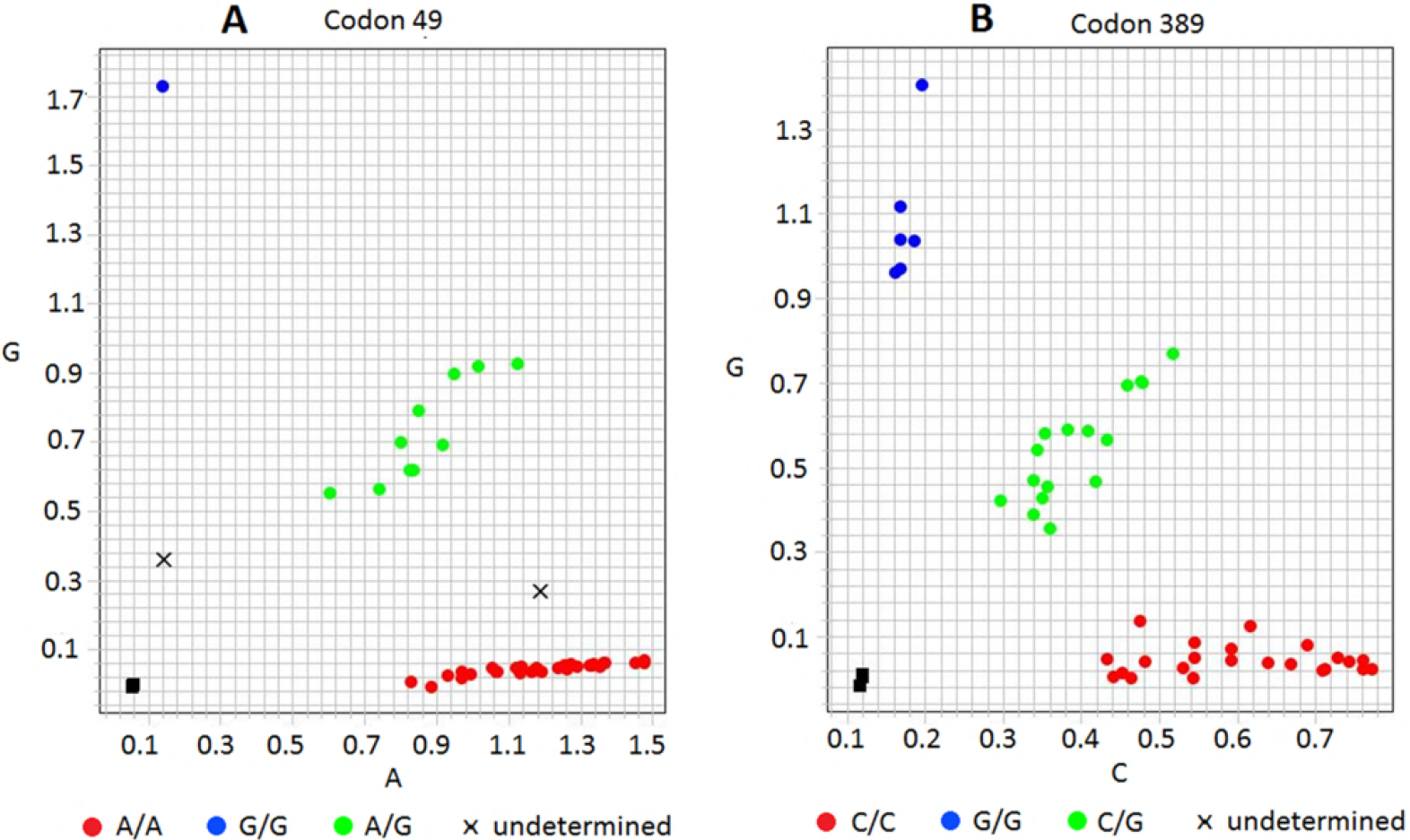
Taqman analysiss of β1-adrenoreceptor gene polymorphism. A: Taqman analysis of Codon 49; B: Taqman analysis of Codon 389.

**Table 1.**
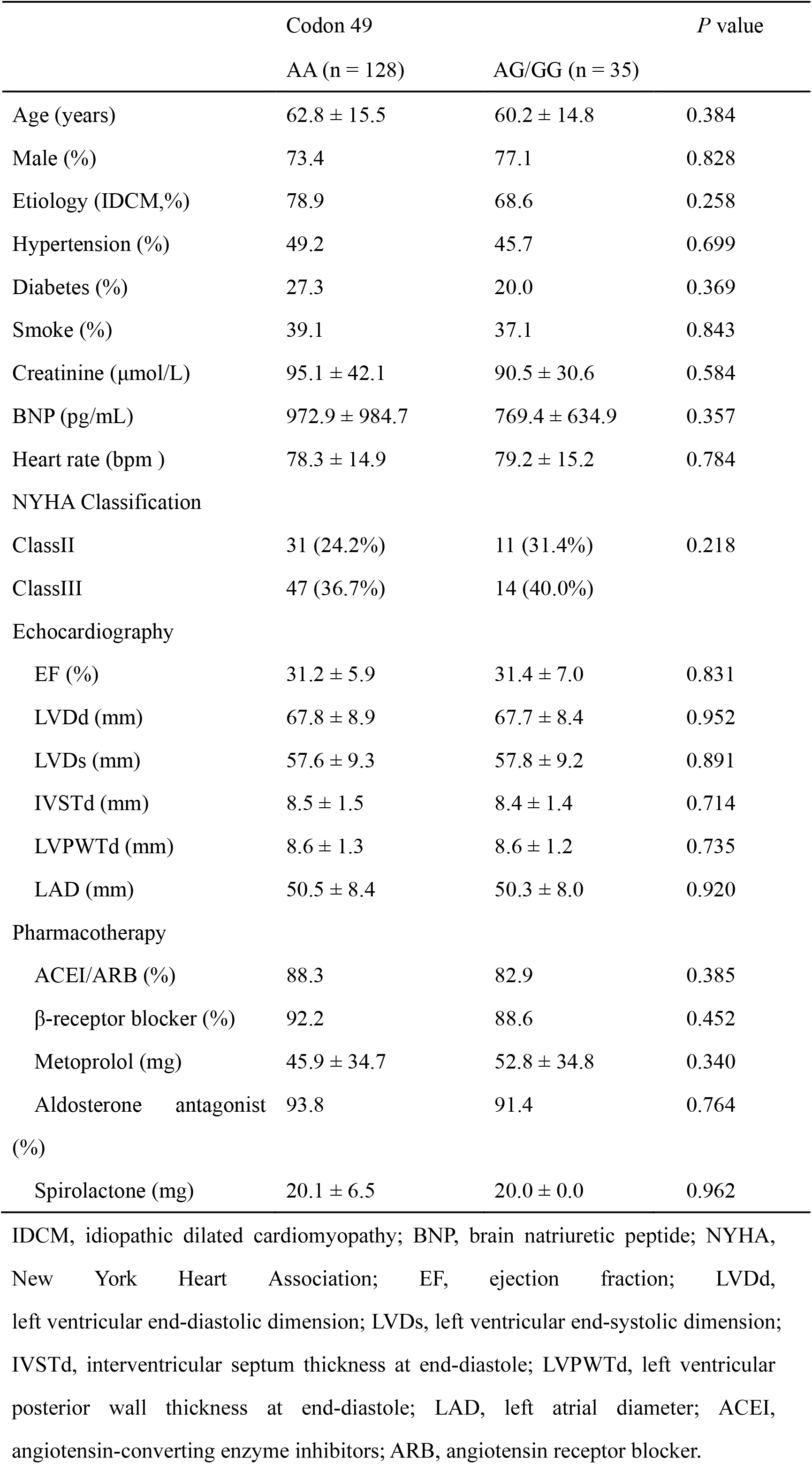
Baseline characteristics. IDCM, idiopathic dilated cardiomyopathy; BNP, brain natriuretic peptide; NYHA, New York Heart Association; EF, ejection fraction; LVDd, left ventricular end-diastolic dimension; LVDs, left ventricular end-systolic dimension; IVSTd, interventricular septum thickness at end-diastole; LVPWTd, left ventricular posterior wall thickness at end-diastole; LAD, left atrial diameter; ACEI, angiotensin-converting enzyme inhibitors; ARB, angiotensin receptor blocker.

|  | Codon 49 |  | *P* value |
| --- | --- | --- | --- |
|  | AA (n = 128) | AG/GG (n = 35) |  |
| Age (years) | 62.8 ± 15.5 | 60.2 ± 14.8 | 0.384 |
| Male (%) | 73.4 | 77.1 | 0.828 |
| Etiology (IDCM,%) | 78.9 | 68.6 | 0.258 |
| Hypertension (%) | 49.2 | 45.7 | 0.699 |
| Diabetes (%) | 27.3 | 20.0 | 0.369 |
| Smoke (%) | 39.1 | 37.1 | 0.843 |
| Creatinine (μmol/L) | 95.1 ± 42.1 | 90.5 ± 30.6 | 0.584 |
| BNP (pg/mL) | 972.9 ± 984.7 | 769.4 ± 634.9 | 0.357 |
| Heart rate (bpm ) | 78.3 ± 14.9 | 79.2 ± 15.2 | 0.784 |
| NYHA Classification |  |  |  |
| ClassII | 31 (24.2%) | 11 (31.4%) | 0.218 |
| ClassIII | 47 (36.7%) | 14 (40.0%) |  |
| Echocardiography |  |  |  |
| EF (%) | 31.2 ± 5.9 | 31.4 ± 7.0 | 0.831 |
| LVDd (mm) | 67.8 ± 8.9 | 67.7 ± 8.4 | 0.952 |
| LVDs (mm) | 57.6 ± 9.3 | 57.8 ± 9.2 | 0.891 |
| IVSTd (mm) | 8.5 ± 1.5 | 8.4 ± 1.4 | 0.714 |
| LVPWTd (mm) | 8.6 ± 1.3 | 8.6 ± 1.2 | 0.735 |
| LAD (mm) | 50.5 ± 8.4 | 50.3 ± 8.0 | 0.920 |
| Pharmacotherapy |  |  |  |
| ACEI/ARB (%) | 88.3 | 82.9 | 0.385 |
| β-receptor blocker (%) | 92.2 | 88.6 | 0.452 |
| Metoprolol (mg) | 45.9 ± 34.7 | 52.8 ± 34.8 | 0.340 |
| Aldosterone antagonist (%) | 93.8 | 91.4 | 0.764 |
| Spirolactone (mg) | 20.1 ± 6.5 | 20.0 ± 0.0 | 0.962 |

**Table 2.** Pharmacotherapy within 1-year follow-up.

|  | Ser49Gly |  | P value |
| --- | --- | --- | --- |
|  | AA (n = 128) | AG/GG (n = 35) |  |
| 3months |  |  |  |
| ACEI/ARB (%) | 92.2% | 91.4% | 0.589 |
| β-receptor blocker (%) | 95.3% | 94.3% | 0.871 |
| Metoprolol (mg) | 50.6 ± 40.7 | 54.2 ± 37.7 | 0.417 |
| Aldosterone antagonist (%) | 94.5% | 91.4% | 0.436 |
| Spirolactone (mg) | 20.2 ± 4.3 | 20.0 ± 0.0 | 0.536 |
| 6months |  |  |  |
| ACEI/ARB (%) | 93.8% | 94.3% | 0.518 |
| β-receptor blocker (%) | 96.9% | 97.1% | 0.389 |
| Metoprolol (mg) | 51.2 ± 38.8 | 53.4 ± 39.2 | 0.875 |
| Aldosterone antagonist (%) | 96.1% | 94.3% | 0.298 |
| Spirolactone (mg) | 20.4 ± 3.1 | 20.2 ± 2.1 | 0.901 |
| 12months |  |  |  |
| ACEI/ARB (%) | 96.1% | 97.1% | 0.629 |
| β-receptor blocker (%) | 97.7% | 97.1% | 0.634 |
| Metoprolol (mg) | 49.8 ± 41.1 | 52.1 ± 40.2 | 0.510 |
| Aldosterone antagonist (%) | 96.1% | 94.3% | 0.398 |
| Spirolactone (mg) | 20.6 ± 2.8 | 20.1 ± 1.8 | 0.878 |

ACEI, angiotensin-converting enzyme inhibitors; ARB, angiotensin receptor blocker.

**Figure 2.**
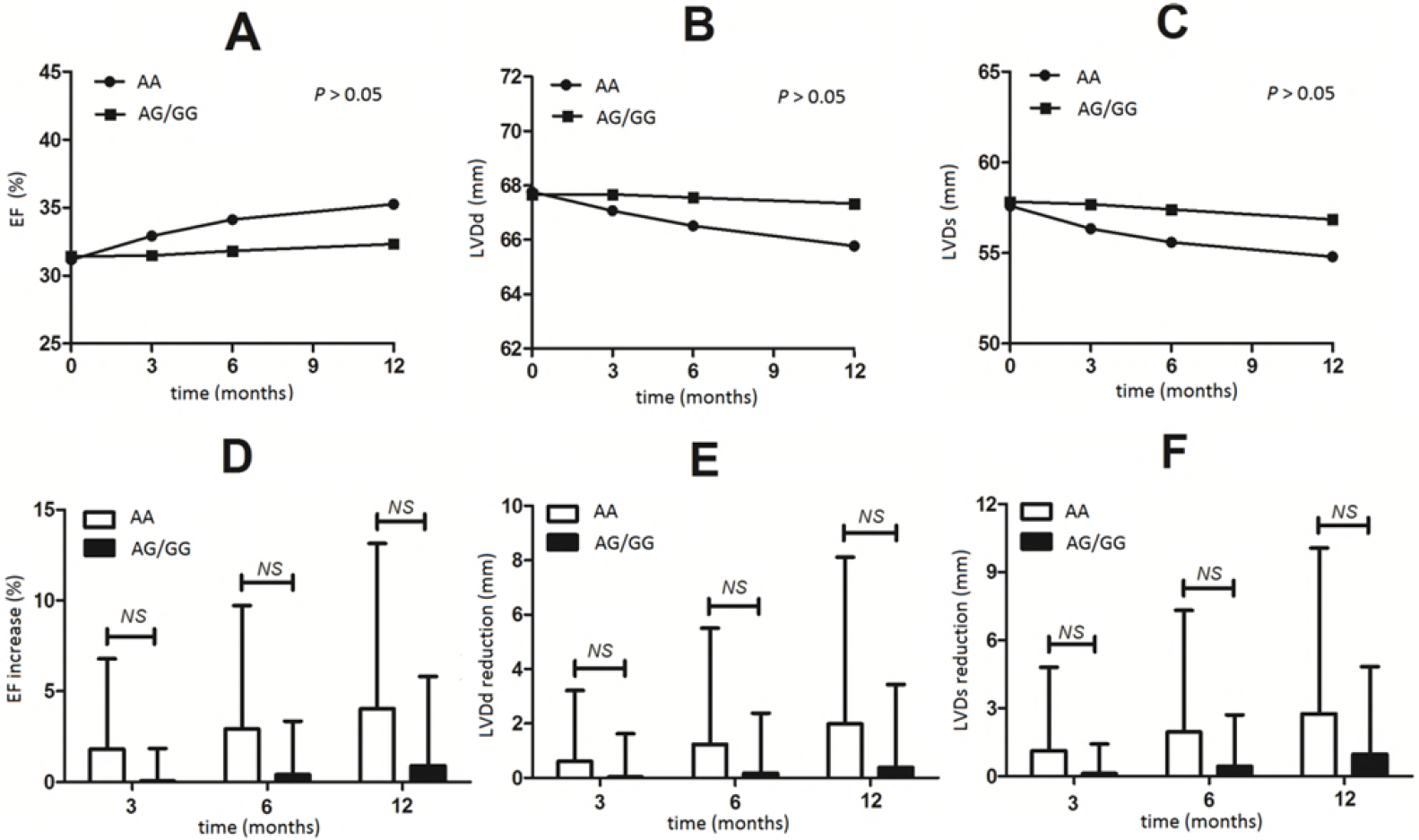
Interaction between Codon 49 genotype of β1-adrenoceptor gene and echocardiographic data. EF, ejection fraction; LVDd, left ventricular end-diastolic dimension; LVDs, left ventricular end-systolic dimension.

**Table 3.**
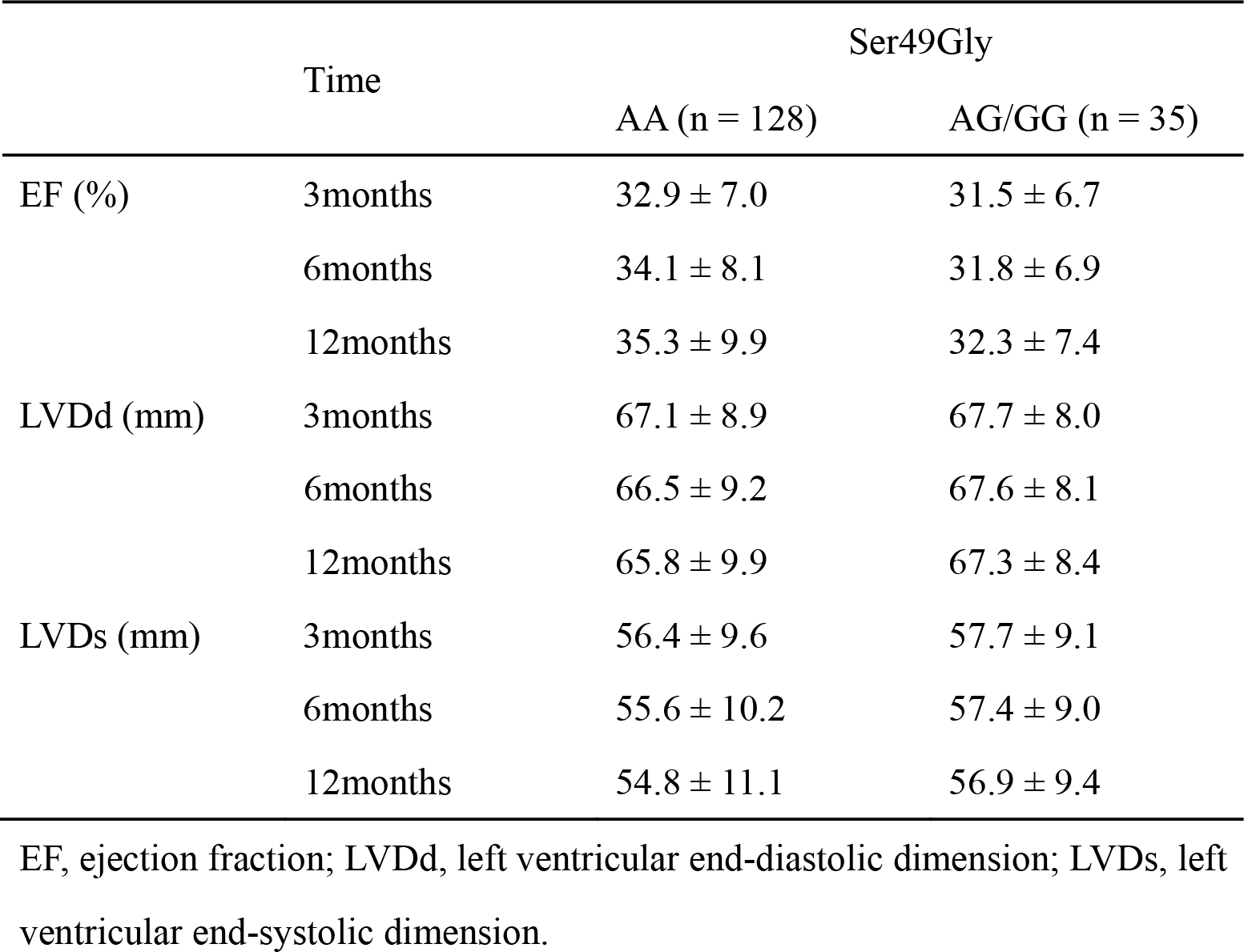
Echocardiography within 1-year follow-up.

|  |  | Ser49Gly |  |
| --- | --- | --- | --- |
|  |  | AA (n = 128) | AG/GG (n = 35) |
| EF (%) | 3months | 32.9 ± 7.0 | 31.5 ± 6.7 |
|  | 6months | 34.1 ± 8.1 | 31.8 ± 6.9 |
|  | 12months | 35.3 ± 9.9 | 32.3 ± 7.4 |
| LVDd (mm) | 3months | 67.1 ± 8.9 | 67.7 ± 8.0 |
|  | 6months | 66.5 ± 9.2 | 67.6 ± 8.1 |
|  | 12months | 65.8 ± 9.9 | 67.3 ± 8.4 |
| LVDs (mm) | 3months | 56.4 ± 9.6 | 57.7 ± 9.1 |
|  | 6months | 55.6 ± 10.2 | 57.4 ± 9.0 |
|  | 12months | 54.8 ± 11.1 | 56.9 ± 9.4 |

EF, ejection fraction; LVDd, left ventricular end-diastolic dimension; LVDs, left ventricular end-systolic dimension.

**Table 4.** Echocardiography within 1-year follow-up.

|  |  | Codon 49 |  |  |
| --- | --- | --- | --- | --- |
|  |  | AA (n = 128) | AG/GG (n = 35) | P value |
| EF increase (%) | 3months | 1.8 ± 5.0 | 0.1 ± 1.8 | 0.078 |
|  | 6months | 2.9 ± 6.8 | 0.4 ± 2.9 | 0.217 |
|  | 12months | 4.0 ± 9.1 | 0.9 ± 4.9 | 0.203 |
| LVDd reduction<br>(mm) | 3months | 0.6 ± 2.6 | 0.1 ± 1.6 | 0.229 |
|  | 6months | 1.2 ± 4.3 | 0.2 ± 2.2 | 0.602 |
|  | 12months | 2.0 ± 6.1 | 0.4 ± 3.1 | 0.461 |
| LVDs reduction<br>(mm) | 3months | 1.1 ± 3.7 | 0.1 ± 1.3 | 0.178 |
|  | 6months | 2.0 ± 5.4 | 0.4 ± 2.3 | 0.430 |
|  | 12months | 2.8 ± 7.3 | 1.0 ± 3.9 | 0.394 |

EF, ejection fraction; LVDd, left ventricular end-diastolic dimension; LVDs, left ventricular end-systolic dimension.

### 3.3. Interaction between Codon 389 genotype and treatment

The analysis results of Codon 389 genotype was shown in figure 1B. There were no significant difference between different genotypes of Codon 389 in baseline characteristics, echocardiography and pharmacotherapy (*P*>0.05, Table 5 and 6). During the whole study, patients carrying G homozygous gene tended to have better EF and left ventricular size improvement compared with C-allele carriers (Figure 3A,3B,3C and Table 7). In further analysis shown in Table 8, Figure 3D,3E and 3F, patients with GG genotype responded better to pharmacotherapy, especially when comparing with those with CC genotype, as shown in EF increase (*P* = 0.000), LVDd reduction (*P* = 0.007) and LVDs reduction (*P* = 0.001), at 12-month time point. Towards the end of the study, similar favorable responses among CG genotype were observed comparing with CC genotype, as shown in EF increase (*P*= 0.019) and LVDs reduction (*P* = 0.020).

**Table 5.** Baseline characteristics.

|  | Codon 389 |  |  | *P* value |
| --- | --- | --- | --- | --- |
|  | CC (n = 79) | CG (n = 69) | GG (n = 15) |  |
| Age (years) | 64.2 ± 15.1 | 61.1 ± 16.0 | 56.7 ± 12.3 | 0.157 |
| Male (%) | 65.8% | 82.6% | 80% | 0.463 |
| Etiology (IDCM ,%) | 77.2% | 76.8% | 73.3% | 0.948 |
| Hypertension (%) | 53.2% | 43.5% | 46.7% | 0.515 |
| Diabetes (%) | 30.4% | 21.7% | 20% | 0.233 |
| Smoke (%) | 46.8% | 29% | 40% | 0.069 |
| Creatinine (umol/L) | 98.2 ± 49.2 | 87.6 ± 25.4 | 102.2 ± 34.8 | 0.253 |
| BNP (pg/mL) | 986.7 ± 1103.1 | 905.8 ± 768.9 | 799.6 ± 711.2 | 0.814 |
| Heart rate (bpm ) | 78.6 ± 15.9 | 79.1 ± 13.7 | 74.9 ± 15.2 | 0.653 |
| NYHA Classification |  |  |  | 0.096 |
| Stage II | 17 (21.5%) | 18 (26.1%) | 6 (40.0%) |  |
| Stage III | 32 (40.5%) | 22 (31.9%) | 7 (46.6%) |  |
| Echocardiography |  |  |  |  |
| EF (%) | 31.7 ± 6.2 | 31.5 ± 5.8 | 27.6 ± 6.4 | 0.051 |
| LVDd (mm) | 68.2 ± 8.4 | 66.8 ± 9.3 | 70.0 ± 7.8 | 0.358 |
| LVDs (mm) | 58.1 ± 8.8 | 56.4 ± 9.8 | 60.9 ± 8.6 | 0.206 |
| IVSTd (mm) | 8.6 ± 1.6 | 8.4 ± 1.3 | 8.3 ± 1.5 | 0.432 |
| LVPWTd (mm) | 8.8 ± 1.4 | 8.4 ± 1.0 | 8.3 ± 1.1 | 0.101 |
| LAD (mm) | 51.0 ± 8.9 | 49.7 ± 7.9 | 50.4 ± 6.5 | 0.644 |
| Pharmacotherapy |  |  |  |  |
| ACEI/ARB (%) | 87.3% | 87% | 86.7% | 0.895 |
| β-receptor blocker (%) | 92.4% | 91.3% | 86.7% | 0.689 |
| Metoprolol (mg) | 44.7 ± 32.0 | 48.2 ± 34.9 | 57.5 ± 47.0 | 0.467 |
| Spirolactone (%) | 93.7% | 94.2% | 86.7% | 0.663 |
| Spirolactone (mg) | 20.3 ± 2.5 | 21.5 ± 8.6 | 20.0 ± 0.0 | 0.479 |

IDCM, idiopathic dilated cardiomyopathy; BNP, brain natriuretic peptide; NYHA, New York Heart Association; EF, ejection fraction; LVDd, left ventricular end-diastolic dimension; LVDs, left ventricular end-systolic dimension; IVSTd, interventricular septum thickness at end-diastole; LVPWTd, left ventricular posterior wall thickness at end-diastole; LAD, left atrial diameter; ACEI, angiotensin-converting enzyme inhibitors; ARB, angiotensin receptor blocker.

**Table 6.** Pharmacotherapy within 1-year follow-up.

|  | Codon 389 |  |  | *P*value |
| --- | --- | --- | --- | --- |
|  | CC (n = 79) | CG (n = 69) | GG (n = 15) |  |
| 3months |  |  |  |  |
| ACEI/ARB (%) | 92.4% | 92.8% | 93.3% | 0.647 |
| β-receptor blocker (%) | 95.0% | 92.8% | 93.3% | 0.712 |
| Metoprolol (mg) | 49.3 ± 34.2 | 51.6 ± 36.6 | 58.2 ± 45.8 | 0.367 |
| Spirolactone (%) | 93.7% | 94.2% | 93.3% | 0.634 |
| Spirolactone (mg) | 20.7 ± 3.0 | 20.8 ± 5.3 | 20.0 ± 0.0 | 0.887 |
| 6months |  |  |  |  |
| ACEI/ARB (%) | 95.0% | 94.2% | 93.3% | 0.299 |
| β-receptor blocker (%) | 97.5% | 97.1% | 100.0% | 0.310 |
| Metoprolol (mg) | 51.2 ± 36.0 | 50.3 ± 37.4 | 56.5 ± 44.2 | 0.276 |
| Spirolactone(%) | 96.2% | 95.7% | 93.3% | 0.421 |
| Spirolactone(mg) | 20.5 ± 1.9 | 20.7 ± 4.8 | 20.0 ± 0.0 | 0.871 |
| 12months |  |  |  |  |
| ACEI/ARB (%) | 96.2% | 97.1% | 93.3% | 0.429 |
| β-receptor blocker (%) | 97.5% | 97.1% | 100.0% | 0.587 |
| Metoprolol (mg) | 50.7 ± 35.8 | 51.6 ± 38.1 | 56.3 ± 42.1 | 0.301 |
| Spirolactone(%) | 95.0% | 95.7% | 93.3% | 0.410 |
| Spirolactone (mg) | 20.6 ± 2.8 | 20.2 ± 1.7 | 20.0 ± 0.0 | 0.892 |

ACEI, angiotensin-converting enzyme inhibitors; ARB, angiotensin receptor blocker.

**Figure 3.**
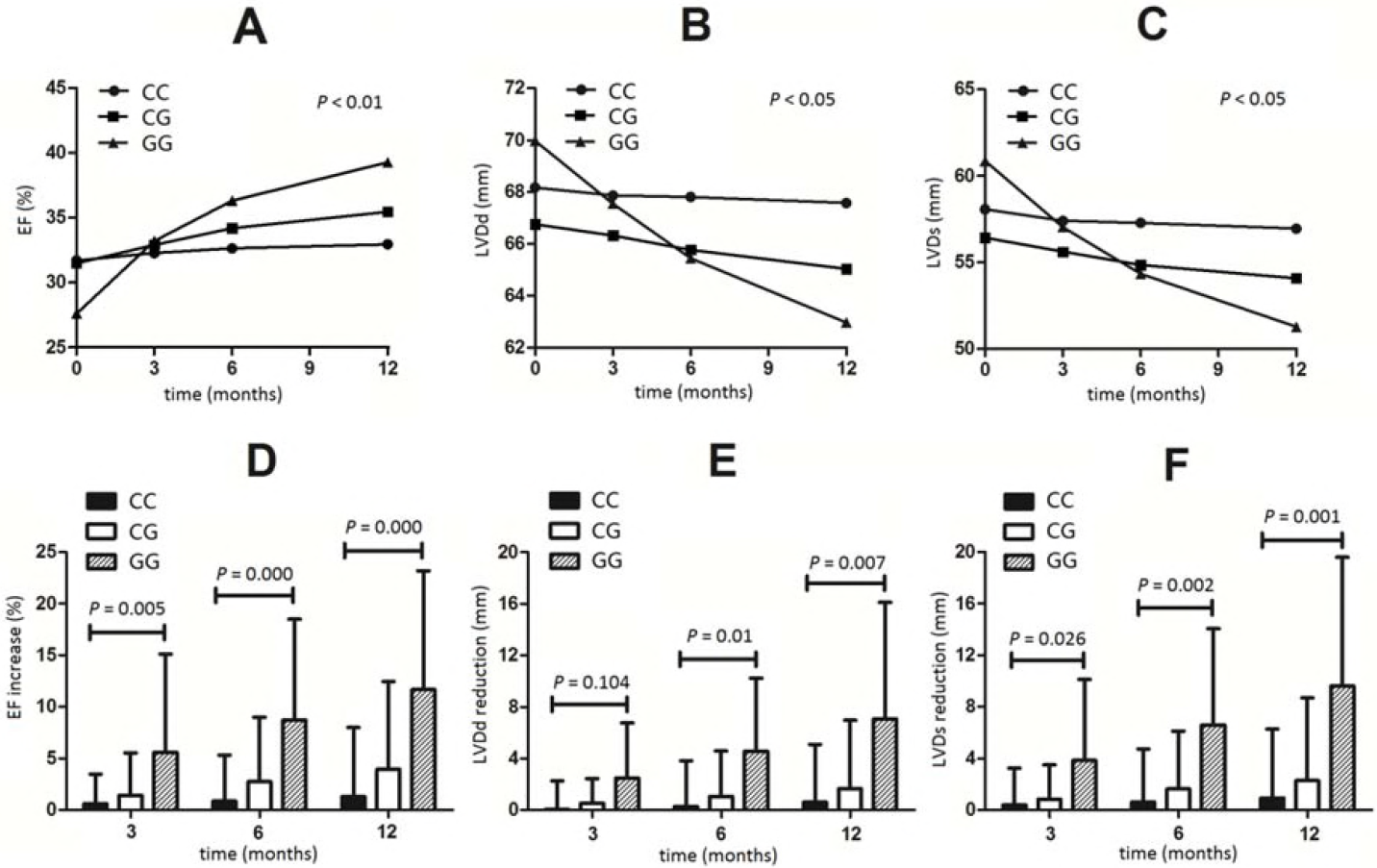
Interaction between Codon 389 genotype of β1-adrenoceptor gene and echocardiographic data. EF, ejection fraction; LVDd, left ventricular end-diastolic dimension; LVDs, left ventricular end-systolic dimension.

**Table 7.** Echocardiography within 1-year follow-up.

|  |  | Codon 389 |  |  |
| --- | --- | --- | --- | --- |
|  | Months | CC (n = 79) | CG (n = 69) | GG (n = 15) |
| EF (%) | 3 | 32.3± 6.5 | 32.9± 6.4 | 33.2± 11.3 |
|  | 6 | 32.6± 7.0 | 34.2± 7.6 | 36.3± 11.9 |
|  | 12 | 33.0± 8.3 | 35.5± 9.5 | 39.3± 13.6 |
| LVDd (mm) | 3 | 67.9± 8.2 | 66.3± 9.1 | 67.6± 9.5 |
|  | 6 | 67.8± 8.4 | 65.8± 9.5 | 65.5± 9.2 |
|  | 12 | 67.6± 8.9 | 65.0± 9.9 | 63.0± 10.6 |
| LVDs (mm) | 3 | 57.4± 8.9 | 55.6± 9.8 | 57.0± 11.3 |
|  | 6 | 57.3± 9.2 | 54.9± 10.3 | 54.3± 12.0 |
|  | 12 | 57.0± 9.9 | 54.1± 10.9 | 51.3± 13.4 |

ACEI, angiotensin-converting enzyme inhibitors; ARB, angiotensin receptor blocker.

**Table 8.**
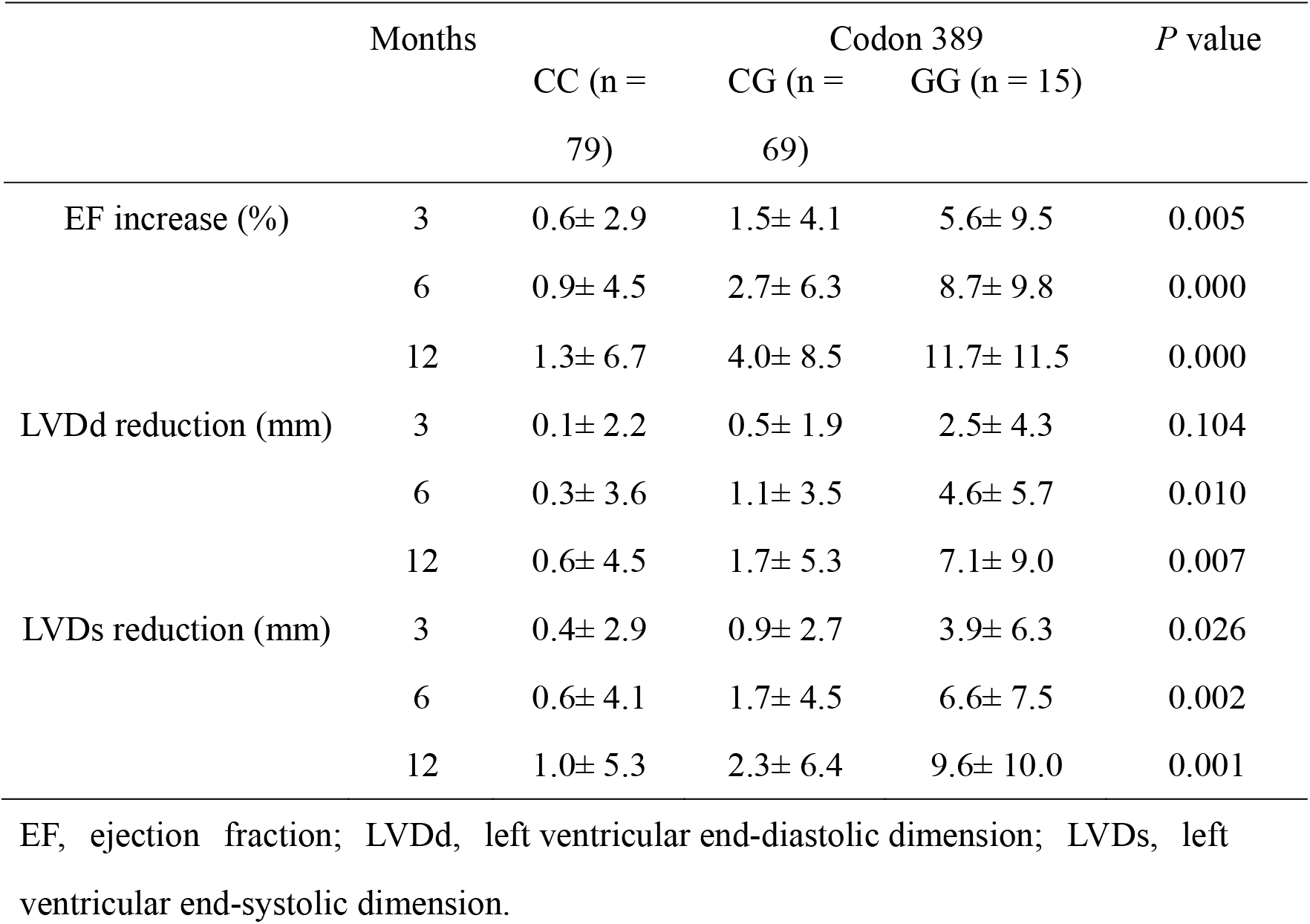
Echocardiography within 1-year follow-up.

As shown in Table 9, participants carrying GG genotype have the lowest MACE incidence (*P*< 0.05) comparing with other genotypes. 1-year MACE incidence among G allele carriers was 9.7%, significantly lower than that among CC genotype (*P* = 0.006).

**Table 9.**
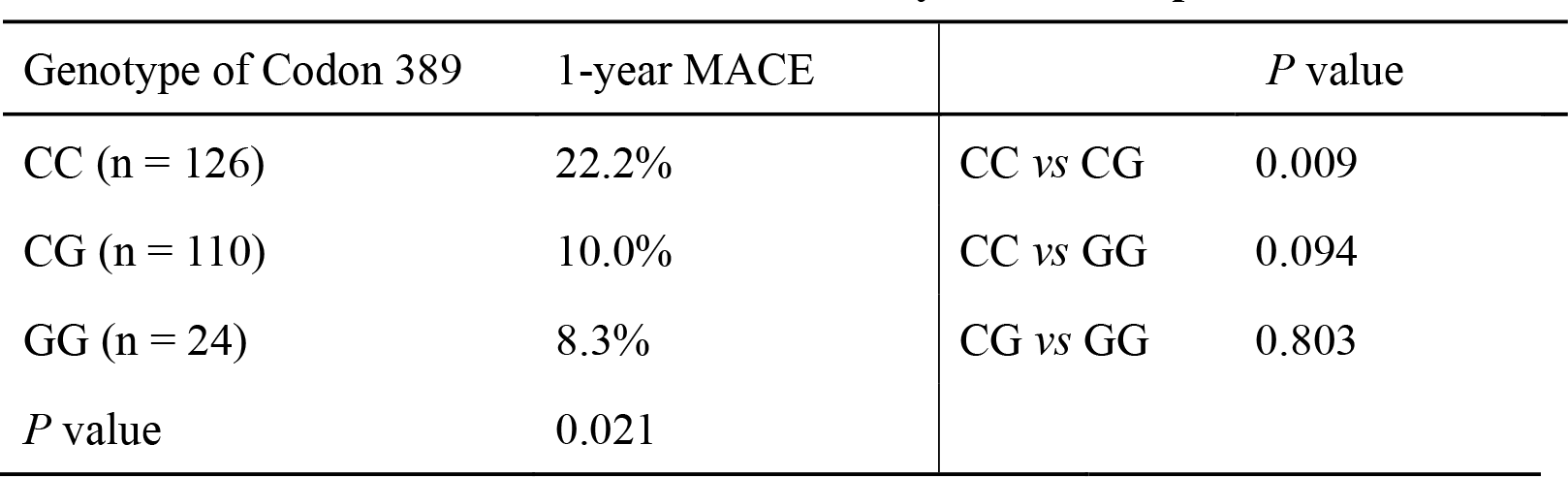
MACE within 1-year follow-up.

| Genotype of Codon 389 | 1-year MACE | *P* value |  |
| --- | --- | --- | --- |
| CC (n = 126) | 22.2% | CC *vs* CG | 0.009 |
| CG (n = 110) | 10.0% | CC *vs* GG | 0.094 |
| GG (n = 24) | 8.3% | CG *vs* GG | 0.803 |
| *P* value | 0.021 |  |  |

## 4. Discussion

Sympathetic-adrenergic system activation is one of the most important compensatory mechanism of CHF which can ameliorative hemodynamic instability, by increasing myocardial contractility and decreasing left ventricular diastolic pressure in the early-phase(1). However, the long-lasting hyperactivation of sympathetic-adrenergic system would induce cardiomyocytes apoptosis, accelerate cardiac remodeling, and down-regulate β-receptor density(1). Substantial researches have demonstrated that β blockers can reduce cardiovascular mortality, incidence of sudden cardiac death and rehospitalization; therefore, the use of β blockers have been recommended as the standard therapeutic medication in all CHF patients by most guidelines(7).

Previous researches demonstrated that gene polymorphisms involved in catecholamine signaling might modulate CHF progression and patients’response to therapy(2); however, the results are often controversial. So there are no definite conclusions to guide clinic applications yet; also, most of these studies were completed among Caucasians or Americans, but rarely Asians(6).

The two primary SNPs of ADRB1 gene are Ser49Gly and Arg389Gly gene polymers. SNP of Arg389Gly in ADRB1 is in intracellular C-terminus, which is an important site for G-protein binding. Arg389, which occurs in 20% - 30% of Asians, has higher basal and isoprenaline-stimulated adenylate cyclase activity, decreased G-protein coupling and reduced β blocker responses. The impact of Arg389Gly on β blocker responses and CHF prognosis were inconsistent in clinical trials. Improvement of cardiac function(4), reduced mortality(5) and higher exercise capacity(8) were observed among CHF patients with Arg389 genotype. However, patients with Gly389X genotype showed greater response to bisoprolol than the Arg389Arg genotype in ABBA study(9), but no response to β blocker treatment in subgroup analysis of MERIT-HF study(10).

Two fundamental researches observed the different influence of Arg389Gly gene polymorphisms of ADRB1 gene on positive inotropic effect of norepinephrine in human atrial myocardium as well: Sandilands et al.(11) found that patients with Arg389 genotype had stronger inotropic effect and cyclicadenosine monophosphate (cAMP) reaction to norepinephrine than other genotypes; while Molenaar et al.(12) did not find such inotropic effects among patients with Arg389 genotype.

In our study, the increase of LVEF, the decrease of left ventricular end-diastolic dimension (LVDd) and left ventricular end-systolic dimension (LVDs) of patients with GG allele was obviously higher than those with CC allele. At 1-year follow-up, the MACE incidence of patients with G allele was obviously lower. From all the above results, we concluded that patients carrying Gly389 gene are more likely to gain more pronounced therapeutic effects of β blocker, better improvement of cardiac function and more promising prognosis.Similarly, another Korean study published in 2016 found that Gly389 carriers weremore sensitive to bisoprolol and had better prognosis comparing with patients with Arg389Arg(9). Biolo A et al. found that Gly389 carriers had better prognosis and higher survival rate in 2008(13) and 2010(14). Also, HF-ACTION DNA substudyshowed that patients with Arg389Arg allele need higher dosage of β blocker than other genotypes in CHF patients(15). These results are all in accordance with our results.

The functional consequences of Gly49 genotype, which occurs inthe N-terminus regionof 14% Asian people, are involved in receptor down-regulation as well as in intracellular trafficking(16). In 2002, Rathz et al.(17) found the affinity to agonist and antagonist of both gene variants were similar; however, Gly49 presented with a more significant receptor down-regulation with long-termagonist treatment compared with Ser49. Also in 2002, Levin MC et al found the adenylate cyclase activity of patients with Gly49 genotype was higher than those with Ser49 while it was more sensitive to inhibitor such as metoprolol(3). The more common Ser49Ser genotype responded less beneficially to β blockers, and this would motivate genotyping to guide doctors to give higher doses for a better clinical outcome(18).

Our study found that there was no statistic difference in cardiac function changes and MACE incidence between different genotypes of Codon 49. It implied that Ser49Gly polymorphism had no obvious influence on therapeutic effects and patients’prognosis, which is in consistence with the conclusion of Humma’s research(19).

This is a retrospective study, so we can not titrage the dosage of β blocker. However, in our study, there is no difference of the dosage of β blocker in patients with different genotypes at 1 year, while the therapeutic effects were obviously worse among patients with CC than those with GG allele, also slightly worse that those with CG allele. We inferred that, if possible, in order to achieve the best therapeutic effect, larger dose of β blocker is required among patients with CC genotype.

Considering CRT-P and CRT-D have some effects on cardiac function of CHF patients, and the treatment of valvular heart disease is mainly depend on surgery instead of drug therapy, our study exclude the above-mentioned cases.

## 5. Conclusions

Our study concluded that Arg389Gly β1-adrenergic receptor polymorphism is related toβ blockers’ therapeutic effect and prognosis of CHF, but not Ser49Gly. With the same dosage of CHF drugs, the improvement of cardiac function and prognosis of patients with Gly389 allele is obviously better than those carrying CC homozygote. This provided a possible gene-related individualized therapeutic plan among CHF patients.

## Acknowledgements

This work was supported by the National Natural Science Foundation of China (grant number 81200092); foundation of the health department of Jiangsu Province (grant number z201514); Nanjing Medical Science and Technique Development Foundation ( grant number ZKX13023).

Author contributions
Rong Gu and Yu Shen contributed this work equally. Lian Wang and Biao Xu are co-corresponding authors. Rong Gu, Yu Shen, Lian Wang and Biao Xu conceived and organized the project and wrote the manuscript. Jianhua Liu and Shuaihui Qiao contributed to experiments and data analysis. All authors discussed the results and commented on the manuscript.

